# Dopamine release effects on striatal blood oxygenation and whole brain plasticity underlying associative learning

**DOI:** 10.1101/2025.07.20.665732

**Authors:** Amir Lawen, Isabella K. Succi, Daniela Lichtman, Darcy S. Peterka, Ishmail J. Abdus-Saboor, Itamar Kahn

**Author notes:** Corresponding Author Columbia University 3227 Broadway Street Jerome L. Greene Science Center New York, NY 10027.

## Abstract

Dopaminergic signaling in the nucleus accumbens (NAc) is central to reward-based learning, but its relationship to brain-wide hemodynamics remains unclear. Using concurrent fMRI and dopamine photometry in awake, behaving mice, we reveal that associative learning induces a gradual temporal shift in NAc blood oxygenation responses that mirrors dopamine release dynamics. This shift emerges with cue-reward learning and extends across a distributed network including prefrontal, insular, and hypothalamic regions. Further, dopamine transients tightly correspond with local BOLD signals, and variations in reward value modulate delayed BOLD responses in both the NAc and additional subcortical structures. Removing dopaminergic contributions abolishes this reward-related modulation, demonstrating that BOLD signals encode dopaminergic value prediction. These findings establish a mechanistic link between dopamine signaling and widespread neural plasticity during learning.

## Introduction

A significant portion of animal behavior can be distilled to the fundamental act of reward-seeking^1^. The ability to form associations between cues and outcomes is critical for enhancing predictability and minimizing harm^2,3^. Classical conditioning, in particular, allows animals to develop motivational states in response to cues that signal potential rewards^4,5^. This learning process engages multiple brain regions, with the nucleus accumbens (NAc) of the ventral striatum acting as a central hub for the acquisition and expression of cue-reward associations^6^. Notably, dopamine release in the NAc is thought to mediate reward-related learning and facilitate motivated responses to incentive cues^7–9^.

While single-cell recordings are essential for identifying regions-specific functions and establishing temporal causality for behavior, they are unable to provide a brain-wide description at the individual level. As brain function depends on activity across multiple brain regions, it is essential to consider whole-brain dynamics to fully understand the neural mechanisms underlying learning^10^. Such an approach can provide a more comprehensive view of brain function and the complex interactions among neural circuits.

Task-based functional magnetic resonance imaging (fMRI) is a well-established technique used in human research for identifying brain regions activated by specific task variables or scheduled events^11^. In humans, the NAc has been implicated in cue-reward learning, reward processing, and encoding reward prediction errors^12–16^. However, little is known about the brain-wide consequences of dopamine release during learning—particularly regarding response plasticity and its contribution to the blood oxygenation level dependent (BOLD) signal changes measured using fMRI in the NAc^12,17,18^.

To address this gap, we used a dual-imaging approach that combines fMRI with dopamine (DA) monitoring in awake, behaving mice engaged in a cue-reward association learning task. Using the GRAB_DA_ fluorescent sensor^19^, we recorded DA dynamics concurrently with fMRI. Our results reveal dynamic shifts in BOLD signal responses that progressively align with reward-predictive cues over the course of learning. Strikingly, similar BOLD dynamics were observed across extensive regions of the striato-pallidal network. Moreover, higher DA release in the NAc was associated with increased local BOLD responses as well as enhanced activity in other key regions, including the anterior cingulate cortex, infralimbic cortex, and amygdala.

## Results

### Associative learning in head-fixed mice during fMRI

To investigate associative learning, we implemented a classical conditioning paradigm. Following habituation, mice were head-fixed but without any additional restraining of the trunk or limb, and a receive-only loop coil was positioned immediately above their head (**Fig. 1A**). In this setup, a blue light served as the conditioned stimulus (CS), followed 1.25 s later by the delivery of a 3 µl water reward (unconditioned stimulus [US]) to water-deprived mice (n = 18). These mice underwent daily sessions over 10 consecutive days, each consisting of 100 trials during which their licking responses were evaluated (**Fig. 1B**). Following familiarization with the licking apparatus, mice performed the task, with their initial exposure to the blue light occurring in the scanner during the first recording session.

**Figure 1.**
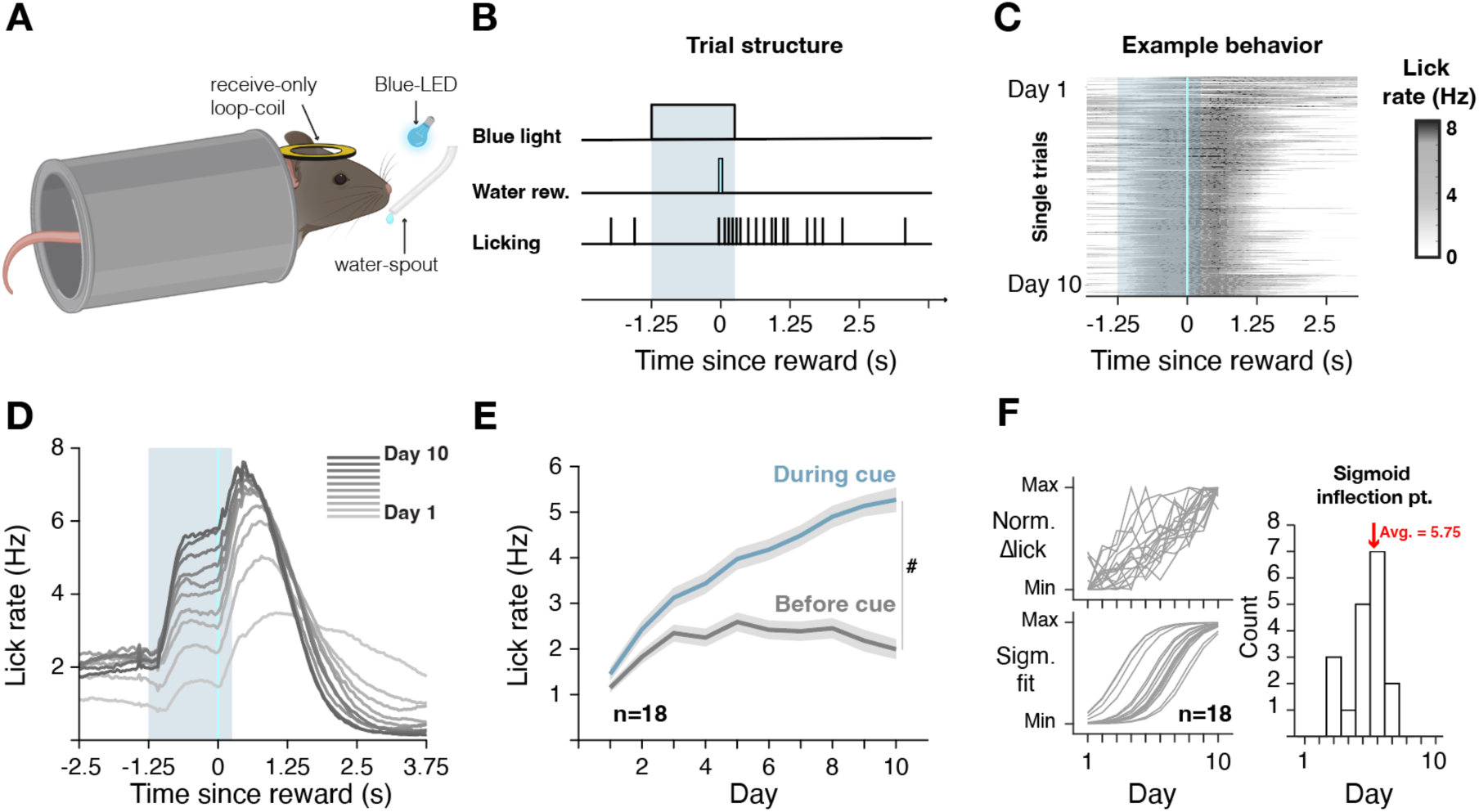
Learning paradigm and mice behavior in the scanner. **A)** Experimental setup depicting of MRI single-element coil positioned above the skull, a waterspout positioned in front of the animal within licking distance and a blue light. **B)** A trial consisted of a blue light turning on for 1.5 s, 1.25 s after the light turned on, a 3 µl water drop was delivered. Individual licks were recorded using video-based analysis. **C)** Lick Behavior from an individual representation animal is shown, demonstrating the emergence of anticipatory licking. **D)** Average group (n = 22) lick rate traces over 10 days. Shading represents blue light period that precedes water delivery. Mice associated the reward with light cue as indicated by the gradual increase in lick rate in the period preceding water delivery. **E)** Anticipatory lick rate during the cued period increased gradually over the course of the experiment while overall baseline (1.25 seconds before cue) lick rate increased as well but plateaued. Average ± SEM. # P<10^-24^. **F)** Individual mouse learning (top, left) shown in a heatmap, with lines representing individual mice anticipatory lick rate difference between the cue period and pre-cue baseline period over the entire period of the experiment. Fitted Sigmoid functions for each mouse (bottom, left) and the distribution of sigmoid inflection points (right) indicate that all animals learned the association during the 10-day period within 5.75 in average (±1.26).ingle-element coil positioned above the skull, a waterspout positioned in front of the animal within licking distance and a blue light. **B)** A trial consisted of a blue light turning on for 1.5 s, 1.25 s after the light turned on, a 3 µl water drop was delivered. Individual licks were recorded using video-based analysis. **C)** Lick Behavior from an individual representation animal is shown, demonstrating the emergence of anticipatory licking. **D)** Average group (n = 22) lick rate traces over 10 days. Shading represents blue light period that precedes water delivery. Mice associated the reward with light cue as indicated by the gradual increase in lick rate in the period preceding water delivery. **E)** Anticipatory lick rate during the cued period increased gradually over the course of the experiment while overall baseline (1.25 seconds before cue) lick rate increased as well but plateaued. Average ± SEM. # P<10^-24^. **F)** Individual mouse learning (top, left) shown in a heatmap, with lines representing individual mice anticipatory lick rate difference between the cue period and pre-cue baseline period over the entire period of the experiment. Fitted Sigmoid functions for each mouse (bottom, left) and the distribution of sigmoid inflection points (right) indicate that all animals learned the association during the 10-day period within 5.75 in average (±1.26).

Associative learning was indicated by the emergence of lick responses that followed the blue light and preceded the water reward (anticipatory licking). Anticipatory lick responses gradually emerged over the course of the experiment (**Fig. 1C, D**). At the population level, there was an increase in anticipatory licking over the learning period, significantly different from baseline lick rates (**Fig. 1E**, F(9,170) = 22.43, *P* < 10^-24^, one-way ANOVA). At the individual level, all mice demonstrated a significant increase in lick rate triggered by the light by the end of the learning phase (**Fig. 1F *top***). A sigmoid fit was used to evaluate the time of learning, with the threshold of learning defined as the sigmoid inflection point. Learning occurred on average at 5.75 ± 1.7 days (mean ± std) at the group level (**Fig. 1F *bottom right*)**.

### Associative learning mediates dynamic transition in BOLD response

Phasic dopamine release in the nucleus accumbens (NAc) has been implicated in signaling reward when it is present prior to learning and in anticipation of it when a stimulus-reward association has been formed^20^. Thus, we first sought to examine NAc response throughout the learning process (**Fig. 2A**). We tracked an indirect measure of neural activity using the fMRI blood oxygenation level-dependent (BOLD) contrast which measures the hemodynamic response. To track brain responses throughout the learning process, we anatomically defined the NAc and extracted the fMRI response. To ensure sufficient signal-to-noise (number of trials) for reliable interpretation, we binned the 10 days into 5 non-overlapping bins. We observed that a response during the delay period emerges during learning (**Fig. 2B**). Due to the relatively long acquisition time for each whole-brain volume with fMRI, a shift in the hemodynamic response function (HRF) can be indirectly inferred by modeling it with a dual gamma function modeling for all data across all days, varying the time-to-peak parameter. We then estimated the change in time-to-peak for each animal across learning bins (**Fig. 2C**). We observed a shift in the best-fit HRF, which was initially characterized by later-rising HRFs averaging roughly 1 s to peak relative to reward delivery, which then transitioned to an earlier-rising HRFs (∼0.2 s to peak, see Methods for details) over the course of learning (**Fig. 2D**; one-tailed student *t*-test: *t*_(17)_ = 4.07, *P* < 0.001). This shift demonstrates a temporal change in the BOLD response, aligning with previous findings in electrophysiological and electrochemical recordings in dopaminergic neurons^21^.

**Figure 2.**
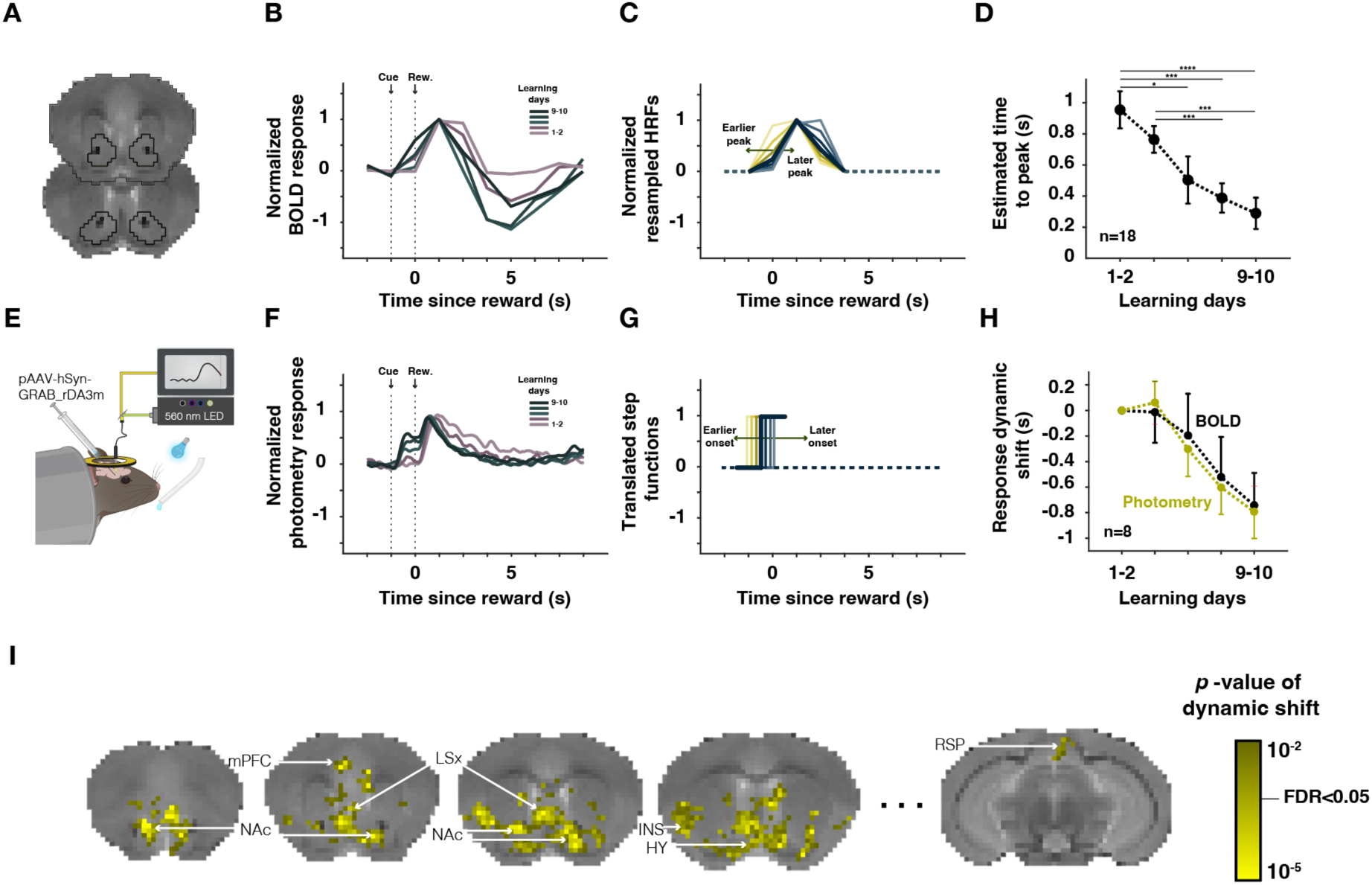
Associative learning mediates dynamic transition in BOLD and dopamine response in the NAc. **A)** Outline of nucleus accumbens overlayed on coronal slices. **B)** Average normalized BOLD traces extracted from nucleus accumbens across the five bins of learning. **C)** Dual-gamma functions used in order to estimate response time, spanning from earlier peak to later peak after cue presentation. **D)** Estimated time to peak of best-fit dual-gamma function, showing a gradual decrease in time to peak. Data are presented as mean ± s.e.m. \**P* < 0.05, \*\**P* < 0.01, \*\*\**P* < 0.005, \*\*\*\**P* < 0.001. **E)** Graphical depiction of dual imaging technique involving fMRI and fiber-photometry, and viral strategy used to label dopaminergic response. **F)** Average normalized dopamine traces (n=8). **G)** Step functions that vary in time-to-onset that were used to estimate response time, spanning from earlier onset to later onset after cue-presentation. **H)** Estimated change of response time compared to first learning bin, both in BOLD signal and GRAB_DA_ photometry signal. Kruskal-Wallis test, *P* = 0.9956. **I)** Dynamic shift across whole brain.

To confirm that the dynamic shift followed dopamine dynamics, we injected a genetically encoded fluorescent sensor, GRAB_DA_^19^, and tracked dopamine release in dopaminergic terminals in the NAc. We estimated the relative fluorescence using fiber-photometry concurrently with fMRI (**Fig. 2E, Supplementary Figure 1**). Using a similar approach as for the BOLD signal, we first estimated time-locked responses to stimulus-reward trials across learning bins and noted a stimulus-related increase in DA (**Fig. 2F**). To quantify this, we estimated response onset by varying a stepwise function onset and finding the best-fit stepwise function with actual fluorescence (**Fig. 2G**). Finally, we quantified the match between the time shift estimated with BOLD and dopamine release measured directly, observing that the fMRI response and GRAB_DA_ signals did not differ (**Fig. 2H**, BOLD compared to Photometry, Kruskal-Wallis test, *P* = 0.9899). The dynamic shift observed in the NAc serves as a robust marker of learning, signifying that the mice successfully associated the cue with the impending reward. To evaluate the brain-wide network involved in reward learning, we hypothesized that regions contributing to this process would exhibit a similar pattern of dynamic shift in their responses. Consequently, we determined the best-fit hemodynamic response function (HRF) for the whole brain across different learning bins (**Fig. 2I**). In addition to bilateral NAc activation, we found that the lateral septal complex, medial prefrontal cortex, insular cortex, and hypothalamus, among other regions, showed significant dynamic shifts across learning bins. This indicates their involvement in orchestrating associative learning.

### FMRI BOLD signal change follows dopamine phasic release in NAc

Dopamine plays a crucial role as a major regulator of medium spiny neurons (MSNs) in the NAc, the primary cell type within this region. However, striatal cholinergic interneurons and other neurotransmitters and neuromodulators also significantly influence cellular activity in the NAc^22–24^. These elements are particularly relevant in regulating striatal dopamine signaling. Given these complexities, it remains unclear to what extent dopamine release contributes to changes in the NAc BOLD signal.

Using a data-driven approach, we aimed to quantify the extent to which the BOLD response mirrored those observed in the GRAB_DA_ signal, evaluating the contribution of dopamine transients to the BOLD response. First, we clustered behaviorally similar trials based on their dopamine transients into four magnitude quartiles (**Fig. 3A**). BOLD responses were then estimated for each dopamine magnitude (**Fig. 3B**). Strikingly, BOLD responses tracked dopamine transients, indicating that dopamine transients are recapitulated in the BOLD signal (**Fig. 3B, C**).

**Figure 3.**
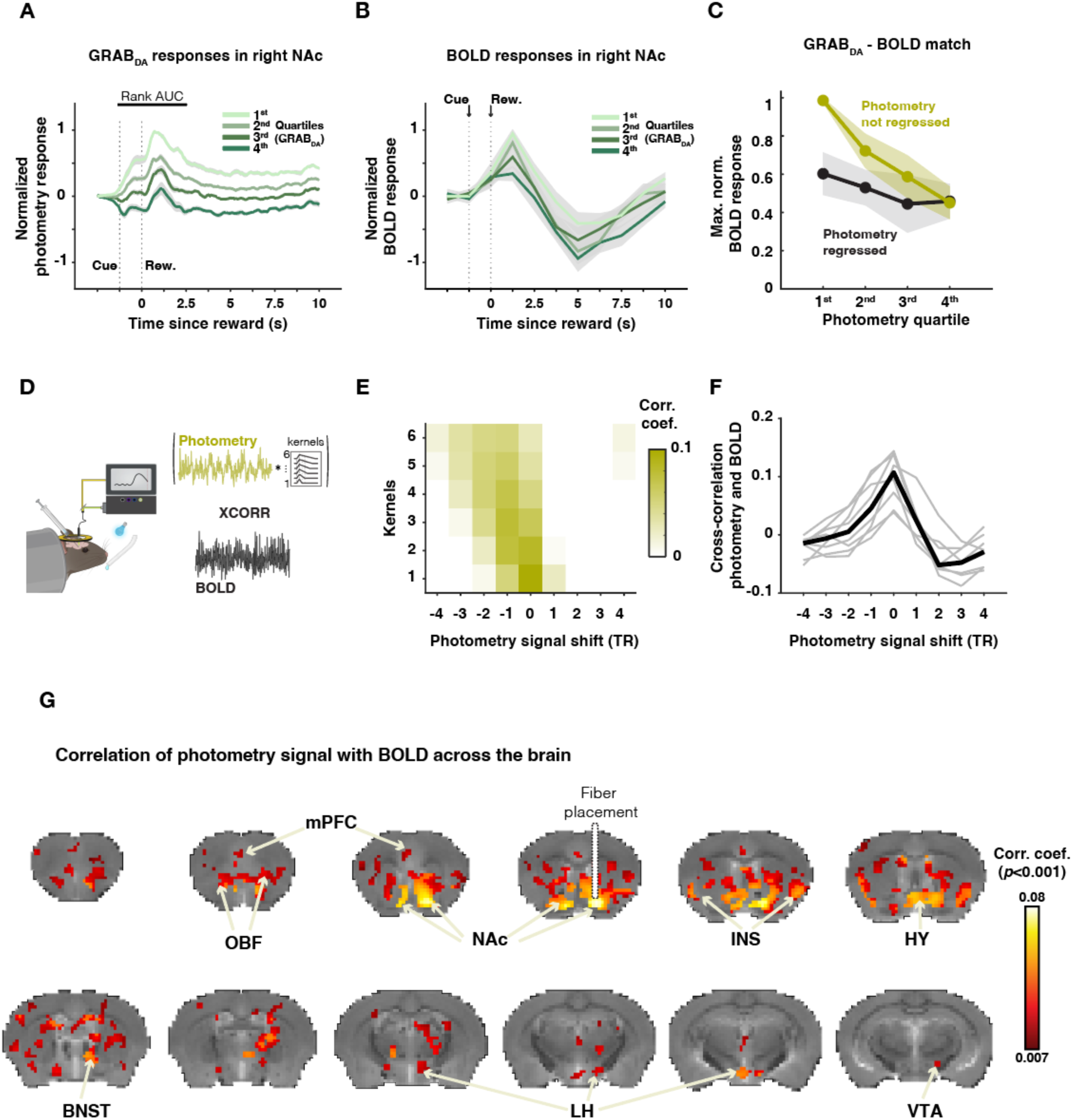
Dopamine phasic release drives BOLD signal change in NAc. **A)** Average photometry response across mice in each photometry response quartile. The rank AUC line indicates the time-window used to calculate the average response for ranking responses into four quartiles. **B)** Average BOLD response in the right NAc across mice in each photometry response quartile. **C)** Quantification of maximal normalized BOLD response across the four quartiles with and without regressing-out the photometry signal. Dots indicate the mean, and shading indicates the s.e.m. **D)** Graphical depiction of the imaging technique and subsequent cross-correlation approach to find the best match between the BOLD and photometry signals. Kernels are the functions used to convolve the photometry signal before cross-correlation. **E)** Heatmap of cross-correlation with different kernels, depicting that the match between BOLD and dopamine release has no lag and a short memory. **F)** Expansion of the first row of (E) to show individual animal compliance. **G)** Upscaled analysis showing the average correlation coefficient between the photometry signal and whole-brain BOLD. Only voxels with a correlation coefficient with P < 0.001 are shown. All colored voxels pass FDR correction.

To determine the temporal transfer function between dopamine transients and the hemodynamic response, we cross-correlated whole-session dopamine signals with the concurrently acquired corresponding BOLD signals in the right NAc (**Fig. 3D**). Dopamine signals were convolved with varying triangular kernels, starting with an identity kernel (Kronecker delta), which has zero memory (Kernel 1), and extending to kernels representing a lasting effect of 6.25 s of decaying effect on future BOLD (representing five time points of BOLD acquisition, Kernel 6). The highest coherence between GRAB_DA_ and BOLD was found with the zero-memory kernel and zero lag, indicating that dopamine release and the BOLD response are co-occurring (up to 0.625 s delay, which corresponds to half of our MRI sequence sampling repetition time; see Methods). This effect was observed at the individual level for each animal participating in this experiment (**Fig. 3F**).

Next, after determining the transfer function between BOLD and GRAB_DA_, we reasoned that upscaling this analysis with zero lag and zero memory to the whole brain would identify regions actively engaged with dopaminergic activity, directly or indirectly, and highlight putative areas relevant for reward processing. This would also help demarcate the regions where dopamine recording has the highest coherence. Indeed, GRAB_DA_ signals showed the highest correlation with the right NAc BOLD signal (proximal to the fiber tip location; **Fig. 3G**). The contralateral NAc was also found to be correlated, suggesting high coherence and equivalent dopamine transients in both NAc regions. Additionally, other regions such as the dorsal striatum, orbitofrontal cortex, insular cortex, thalamus, and ventral tegmental area (VTA) displayed reliable correlations, underscoring their important contributions to the stimulus-reward learning task.

### Reward value representation in NAc BOLD reflects dopaminergic release

Beyond its role in associating stimulus and expected reward, the nucleus accumbens (NAc) is characterized as a center for reward processing, valence, and value representation across species ^25,26^. The major neurotransmitter implicated in these processes in the NAc is dopamine, which is released from mesolimbic axons originating in the VTA or substantia nigra pars compacta. It has been previously shown that optogenetic stimulation of dopaminergic neurons in the midbrain results in a positive BOLD response in the ventral striatum ^27,28^. However, knowledge about the correspondence between dopaminergic release transients related to received reward (rather than anticipated reward as explored above) and the BOLD signal during voluntary behavior is lacking.

To address this, mice participated in a second experiment, where the trial temporal structure remained the same, but reward volumes were varied randomly between 1, 3, and 6 µl (**Fig. 4A, B**) maintaining the predictive nature of the light cue that a reward is expected, but varying the magnitude such that volume might be similar to the volume they previously experience, lower or greater. Indeed, while mice showed anticipatory licking responses for the upcoming reward, as established before, these responses did not differ as a function of reward volume (**Fig. 4C** Nevertheless, once reward was dispensed, robust lick rate gradation for reward magnitude was observed (**Fig. 4D**).

**Figure 4.**
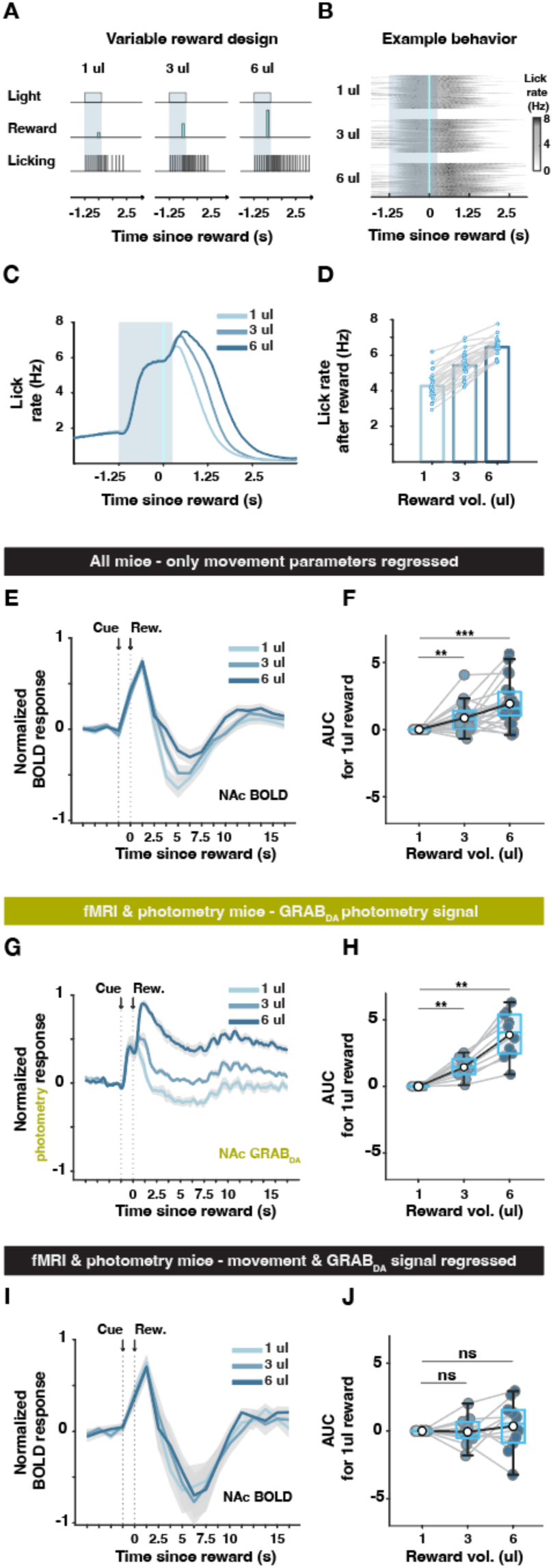
Reward value representation in NAc BOLD reflects dopaminergic release. **A)** Similar to trials during the learning phase, in the second experiment, a trial consisted of a blue light turning on for 1.5 s. 1.25 s after the light turned on, a water drop was delivered, with the reward volume randomly varied between 1, 3, and 6 µl. Individual licks were recorded using video-based analysis. **B)** Lick behavior from a representative animal is shown, demonstrating similar behavior before reward delivery and then the expected divergence in behavior during reward consumption. **C)** Average group (n = 22) lick rate traces over 5-7 days. Shading represents the blue light period that precedes water delivery. Mice display similar behavior across all trial types prior to reward delivery, followed by a divergence in behavior corresponding to the reward volume. **D)** Average lick rate quantification in the 2.5 s window post-reward delivery. **E)** BOLD response extracted from the right-side nucleus accumbens for the three trial types. The line represents the average across animals, and shading represents the standard error of the mean (s.e.m.). **F)** Quantification of the area under the curve compared to trials with a 1 µl reward volume over 12.5 s (10 fMRI time-points). N = 22. ** P < 0.01, *** P < 0.005. **G)** Average GRAB_DA_ photometry time course from the right NAc during the three trial types. Gray shading indicates s.e.m. **H)** Same as in (F) but for the photometry signal. ** P < 0.01. **I)** BOLD response extracted from the right NAc for mice with concurrent fMRI and photometry recordings, with the photometry response regressed out from the BOLD signal. **J)** Same quantification as in (F) and (H).

NAc BOLD time courses showcased an identical response across all trial types from cue onset and up to 2.5 s (**Fig. 4E**), potentially reflecting the response to the cue prior to reward presentation. Reward magnitude-related signals become observable 2.5 s after reward delivery. The lower the reward, the more extreme the undershoot below baseline (**Fig. 4E, F**; two-tailed Student *t*-test: 6 µl compared to 1 µl, t(21) = 4.5, *P* = 9.14×10^-5^; 3 µl compared to 1 µl, t(21) = 3.44, *P* = 0.0012; 6 µl compared to 3 µl, t(21) = 2.61, *P* = 0.0081).

To determine whether the late gradation in the BOLD response is correlated with dopaminergic release, we again acquired fMRI and GRAB_DA_ fiber photometry concurrently. As expected, the GRAB_DA_ signal had an identical component across trial types preceding reward delivery. However, following reward delivery, all trial types exhibited a swift increase in GRAB_DA_ signal, which then decayed in order from 1 µl to 6 µl. Quantifying the area under the curve similarly to the previous BOLD analysis resulted in a similar gradation (**Fig. 4G, H**; two-tailed Wilcoxon signed-rank test: 6 µl compared to 1 µl, *P* = 0.002; 3 µl compared to 1 µl, *P* = 0.002; 6 µl compared to 3 µl, *P* = 0.002; n = 10).

Finally, for a subset of mice that underwent the dual imaging approach, we evaluated the contribution of dopamine in NAc by regressing out the concurrent GRAB_DA_ signal from the BOLD signal and then estimated the resultant BOLD signal. Remarkably, the late signal gradation was eliminated (**Fig. 4I, J**; two-tailed Wilcoxon signed-rank test: 6 µl compared to 1 µl, *P* = 0.6250; 3 µl compared to 1 µl, *P* = 0.8457; 6 µl compared to 3 µl, *P* = 0.4922; n = 10), establishing a clear and novel link between BOLD reward representation in the NAc and dopaminergic release.

### Brain-wide reward value representation

After establishing that the later response in NAc BOLD reliably represents value, we reasoned that we could perform this analysis across the whole brain to identify additional regions that encode value. We defined a general linear model (GLM) in which we fit two separate, non-overlapping HRFs representing the cue- and reward-related responses separately, with the reward-related response parametrically modulated by reward volume (1, 3, and 6 µL). We then computed a brain-wide map of voxels whose late response was significantly modulated by reward volume (**Fig. 5**).

**Figure 5.**
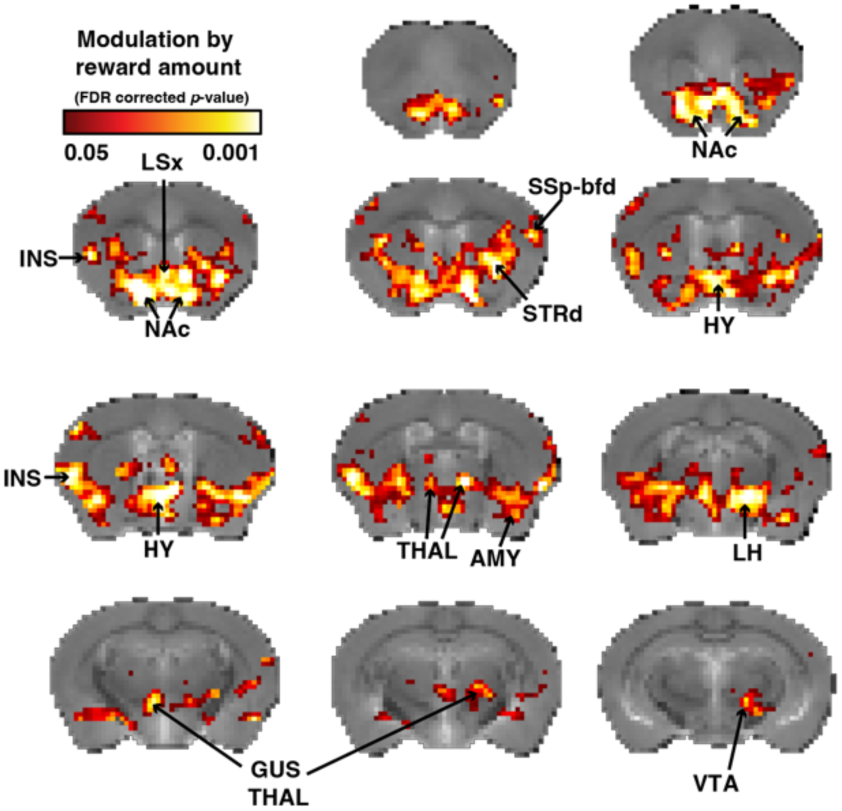
Whole brain representation of value. Maps representing statistical significance of positive parametric modulation of the late response. NAc – Nucleus accumbens, INS – Insula, LSx – Lateral septal complex, STRd – Dorsal striatum, SSp-bfd – Primary somatosensory cortex; barrel field, HY – Hypothalamus, THAL – Thalamus, AMY – Amygdala, LH – Lateral hypothalamus, GUS THAL – Gustatory thalamus, VTA – ventral tegmental area.

The maps showed a significant symmetric parametric modulation in the NAc, as expected. Beyond the NAc, other regions that encoded value included the lateral septal complex (LSx; previously shown to exhibit dynamic shift and learning-related modulation), primary somatosensory barrel field cortex (SSp-bfd), dorsolateral striatum (STRd), anterior and posterior insula (INS), hypothalamus (HY), amygdala (AMY), thalamus (THAL), gustatory thalamus (GUS THAL), lateral hypothalamus (LH), and the ventral tegmental area (VTA).

## Discussion

The ability to associate a cue with an upcoming reward is an adaptive behavior that enables animals to allocate attention and resources more effectively. However, tracking the neural processes that underlie learning over prolonged periods of time remains challenging. To our knowledge, this study is the first to track brain-wide indirect measures of neural responses in parallel with dopamine-specific responses in the ventral striatum during associative learning. We demonstrate that associative learning is correlated with shifts in BOLD responses in the nucleus accumbens (NAc), which reliably correspond with GRAB_DA_ photometry signals. These shifts are not limited to the NAc; the lateral septum (LSc) and medial prefrontal cortex (mPFC) also show prominent changes. Although dynamic shifts in dopaminergic release and neural activity during associative learning have been reported in the NAc^21,29–31^, prefrontal cortex^32^, and septal complex^33^, our findings are the first to offer a non-biased, concurrent, and spatially localized characterization of the broader network correlated with learning. Notably, our data also implicate the insular cortex and hypothalamus–regions not previously established in this context.

That said, our analysis may not capture all regions involved in learning. Some regions may exhibit hemodynamic response functions (HRFs) we did not model, may not produce a reliable hemodynamic response, or suffer from low signal-to-noise, especially in midbrain structures. Additionally, regions more susceptible to motion artifacts—such as the midbrain, pons or cerebellum—may not be accurately represented.

A key contribution of our study is clarifying how dopamine influences brain-wide BOLD signals. This question has not been explored in a minimally invasive manner in awake, behaving animals. Prior work has suggested mixed effects of dopamine on the striatal BOLD response^28,34,35^. For example, stimulating D1 or D2 medium spiny neurons (MSNs) in the striatum produces distinct BOLD profiles—positive or negative, respectively^35^—while different input nuclei elicit varied responses^28^. Li and Jasanoff^33^ showed that inhibition of D1 or D2 receptors did not significantly alter BOLD amplitude, supporting the hypothesis that dopamine may affect BOLD signals via direct vasoactive mechanisms rather than through post-synaptic metabolic demand. Yet, they also observed widespread BOLD changes after receptor inhibition, suggesting dopamine’s role in modulating neural firing. These complexities make interpreting the BOLD signal challenging. Our findings provide foundational evidence that dopamine plays a selective role in driving the BOLD response. We observed a correlation between dopaminergic release and BOLD signals (**Figure 3C**), supporting the idea that dopamine may result in vasodilation either directly or indirectly. Given that only supraphysiological dopamine release reliably alters NAc spiking activity^36^, and that our results are derived from naturalistic conditions, this favors a direct vasoactive role. Still, Li and Jasanoff’s findings suggest dopaminergic effects on neuronal firing even when BOLD amplitude changes are minimal. Thus, we cannot conclusively rule out either mechanism and instead propose a hybrid model. This is supported by our finding that the lowest quartile of GRAB_DA_ signals does not impact BOLD when regressed out (**Figure 3C**), indicating dopamine-independent contributions—possible from cortico-striatal input to D1 or D2 MSNs. These inputs have been shown to elicit positive BOLD responses and are essential for driving motivated behavior^35,37–39^. Moreover, regressing out the dopamine signal did not eliminate the BOLD dynamic shift in the NAc (**Fig. 2H**), further supporting a role for cortico-striatal inputs in shaping this response.

The initial BOLD response in the NAc appears to be largely driven by phasic dopamine release (**Fig. 3C**). However, dopaminergic reward prediction error signals—and corresponding BOLD responses—include two distinct components following learning. As Schultz et al. (2016) noted, dopamine signaling evolves from a broad, unselective response to a more refined signal reflecting subjective value and marginal utility^40,41^.

We observed this two-phase pattern during the variable reward phase: an initial general response followed by a graded signal tracking reward value. This delayed, value-sensitive BOLD component could be attributed to dopamine signaling, as regressing out GRAB_DA_ eliminated value-dependent differences (**Fig. 4I, J**). This enabled us to identify value-sensitive regions throughout the brain (**Fig. 5**), including the basal ganglia, thalamus, hypothalamus, and other subcortical regions. These findings align with prior studies showing that value encoding emerges in the cingulate cortex during early learning but diminishes with task proficiency^42^. Since variable rewards were introduced only after mice had learned the task, we did not observe value encoding in high-order cortices.

Finally, the infralimbic cortex—a subregion of the mPFC critical for controlling motivated behaviors and participating in decisions for action execution^32,43,44^—emerged as central in our analyses. It exhibited BOLD dynamic shift (**Fig. 2I**) and coherence with dopaminergic signals (**Fig. 3G**). These findings reinforce its established role in guiding motivated behavior and associative learning, particularly through projects that facilitate conditioned reward responses^32,45^.

## Methods

### Animals

All procedures were conducted in accordance with the ethical guidelines of the National Institutes of Health and were approved by the Institutional Animal Care and Use Committee (IACUC) at Columbia University. Eighteen male C57/BL6 mice (2–3 months old) were implanted with MRI-compatible head posts and housed in a reversed 12-hour light/dark cycle. After a recovery period of 7–10 days, during which the mice returned to their preoperative weight, a 7–10 day water restriction regime began. The mice were initially given water access for 120 s per day, gradually decreasing to 90 s until they stabilized at approximately 85% of their initial weight. Additional water access time was provided if the mice did not meet the 85% weight criteria. Once weight stabilization was achieved, the scanning sessions commenced.

### Head-post surgery

For head fixation, mice were implanted with custom-made MRI-compatible head posts. Pre-operatively, the mice received extended-release Buprenorphine (Buprenorphine XR) for analgesia. They were then anesthetized with isoflurane (1.5–2.5%) and given additional local analgesia (Marcaine) before making any incisions. The scalp and periosteum layer were removed to expose the skull, and the head post was secured using dental cement (Metabond, Parkell). In some cases, an additional layer of dental cement (Paladur, Heraeus Kulzer) was applied to enhance homogeneity and reduce susceptibility artifacts.

### Head fixation and scanner habituation

After recovery, mice were handled daily for progressively longer periods. Initially, the mice were allowed to sniff the experimenter’s hand, followed by gradually lifting them and letting them explore the experimenter. Subsequently, the mice were introduced to the cradle used for head fixation. After two days of handling, the mice were allowed to enter the head-fixation cylinder. Once inside, the experimenter captured and fixed the head post with fixation forks. The mice were head-fixed for progressively longer durations—1, 2, 5, 8, 15, and finally 25 minutes on consecutive days. During the last two days of habituation, the mice were placed inside the magnet bore, where scanner noises were introduced.

### Water delivery

Water was delivered using a normally closed Parker Series 3 Miniature Inert Liquid Valve, which was triggered for a preset and calibrated amount of time to deliver 1, 3, or 6 µl of water per trial. Since the valve is not MRI-safe, it was placed outside the MRI room and connected to the experimental setup in the scanner via a 7-meter-long tube. At the start of each scanning day, water was flushed through the tube for 4 minutes to ensure the dissipation of air bubbles. Water was then calibrated by administering 100 drops into a small container, which was weighed to calculate the average water drop volume. Calibration was iteratively adjusted until accurate volumes were achieved.

### Behavioral pretraining

Although the mice were entirely naive to the structure of the experiment and were introduced to the unconditioned stimulus only on the first day of recording, it was crucial to familiarize them with the setup, particularly the water delivery system. On the last day of habituation, at random time points, the mice were given 3 µl water drops from the water spout positioned near their mouths. We ensured that they licked the water drops. If a mouse did not lick, the experimenter gently moved the water spout to touch the mouse’s mouth, prompting the water-deprived mouse to initiate licking. By the end of the last day of habituation, each mouse voluntarily acquired 10 drops of water.

### Behavioral monitoring

To monitor mice and extract behavioral features, the setup included an MRI-compatible 12M-i camera (MRC Systems GmbH, Heidelberg, Germany) with an acquisition rate of 29.97 frames per second and a built-in tunable infrared LED to optimize brightness and contrast. The scanner bore is small, preventing the camera from capturing a side view of the mouse. Importantly, tracking tongue protrusion is challenging from a front view since the tongue lacks proper contrast in a grayscale image from that perspective. Therefore, we incorporated a mirror positioned by the mouse’s side and tilted at 45°, allowing the front-view camera to also capture a side view.

To detect licks, we used an in-house algorithm. Briefly, we performed region of interest (ROI) analysis on the tip of the spout. With the grayscale camera, infrared LED, and black resin setup, the tongue appears bright white on a dark background once protruded. Therefore, when the tongue approaches the designated ROI, the overall brightness of the ROI increases reliably. After estimating the brightness of the ROI across a run, a simple spike-detection approach is employed.

### Image acquisition

MRI scans were performed at 9.4 Tesla MRI (Bruker BioSpin, Ettlingen, Germany) using a quadrature 86 mm transmit-only coil (Bruker BioSpin) and a 20 mm loop receive-only coil (Bruker BioSpin). Mice were briefly anesthetized (5% isoflurane) and mounted on the cradle. Water port was adjusted until proper access to water reward was achieved. Each daily session included acquisition of one low-resolution rapid acquisition process with a relaxation enhancement (RARE) T2-weighted structural volume (50 coronal slices, TR/TE 2300/8.5 ms, RARE factor = 4, flip angle = 180°, 200 × 200 × 300 μm^3^, field of view of 19.2 × 19.2 mm^2^, matrix size of 96 × 96) and multiple six minutes spin-echo echo-planar imaging runs measuring BOLD (TR/TE 2500/13.022 ms, flip angle = 90°, 50 coronal slices, 200 ×200× 300 μm^3^, field of view of 14.4 × 9.6 mm^2^, matrix size of 72 × 48; the imaged volume was framed with four saturation slices to avoid wraparound artifacts). During scanner calibrations and acquisition of anatomical images, mice received random drops of water to keep them calm.

### Data pre-processing and confound correction

Scans were organized according to BIDS format ^46^. First, raw data underwent noise reduction using Noise reduction with Distribution corrected (NORDIC) PCA ^47^. Pre-processing was performed on separate runs using Rodent Automated Bold Improvement of EPI Sequences – RABIES ^48^. Pre-processing included inhomogeneity correction, motion correction, a rigid registration between functional and anatomical scan per session, non-linear registration between a mouse scanning session anatomical scan and provided RABIES template, a common-space re-sampling to 0.2 x 0.2 x 0.45 mm^3^. After pre-processing, a visual inspection was performed on all data for quality control and session exclusion. Later, pre-processed data was confound-corrected using ICA-AROMA^49^, white matter; ventricle; and global signal regression, motion regression, spatial smoothing to 0.5 mm^3^.

### Determining HRF and shift in HRF response

After confirming the mice’s ability to associate the light stimulus with the water reward learning in the scanner, we sought to identify brain regions subserving learning by using fMRI blood oxygenation level-dependent (BOLD) contrast. Prior research highlighted differences between the mouse and human hemodynamic response function (HRF), primarily in response duration. Consequently, using the human canonical HRF would not accurately capture the true responses within the mouse brain. The mouse HRF, characterized by shorter responses, combined with constraints in increasing the sampling rate due to signal-to-noise issues and relaxation times, results in relatively under-sampled data compared to human studies. These constraints magnify the impact of minor deviations on analysis efficiency significantly. Moreover, studies have shown that the HRF is notably influenced by brain states, affecting factors such as time-to-onset, magnitude, and dispersion. This adds another layer of uncertainty in accurately characterizing the mouse HRF. Given these factors, it was necessary to comprehensively and reliably characterize task-evoked BOLD responses, considering the unique nuances of the mouse HRF. Therefore, average BOLD response from all runs across all animals was calculated. Afterwards, using SPM’s function, *spm_hrf*, we fit the BOLD response and found the specific parameters to best fit the response. Consecutively, the time-to-peak parameter in *spm_hrf* function was gradually tuned to find best-fit-hrf parameters per mouse per learning bin. These parameters were later used for further GLM whole-brain response HRF fitting.

### Functional connectivity analysis

For functional connectivity analysis, we computed beta-series correlations. Single trial betas were estimated using a general linear model (GLM), where each column contained only one trial. A beta-series was estimated for each voxel. Seed’s beta series was estimated as the median beta series across all included voxels in the seed. After estimating betas for each voxel or seed, pairwise Pearson correlation was computed per animal. Pearson’s Rho value per animal were then Fisher’s-Z transformed, and transformed values were used for second level statistical inference across animals.

### Fiber photometry

Ceramic sleeves (Thor Labs) were used to connect the implanted ceramic ferrule and fiber implant to the MRI-compatible patch cord (custom-made by DORIC). Signals were recorded in real time using a real-time processor (RZ10X, TDT) and extracted with Synapse software (TDT). A 465 nm LED was used to excite GRAB_DA_, while a 405 nm LED served as an isosbestic control to account for fluorescence changes caused by movement artifacts and photobleaching.

To maintain signal quality and magnitude, a dummy recording was initiated during positioning in the MRI bore and then halted during MRI adjustments, shimming, and anatomical scan acquisition to prevent unnecessary photobleaching. A custom MATLAB script synchronized the MRI scan with the photometry recording using an analog trigger output from the MRI scanner. The script also executed the task design, continuously prompting Synapse to maintain temporal alignment and correct for temporal drift.

### Fiber photometry analysis

TDT data folders were directly imported into custom MATLAB analysis scripts. To ensure smooth physiological trends, raw time-series data from each session were filtered using a median filter. To correct for magnitude drift due to photobleaching, first-order linear trends were removed over 10-minute scanning sessions while preserving the average signal magnitude (detrending without demeaning).

The raw signal was then downsampled from ∼1 kHz to 50 Hz, as the original sampling rate was unnecessary given the slow kinetics of the signal (τ-on > 0.5 sec in the NAc in vivo). This reduction minimized data size and optimized processing efficiency. ΔF/F was calculated using a dynamic baseline, determined for each time point by averaging the preceding 64 seconds. The resulting ΔF/F signal was then smoothed using robust locally weighted scatter plot smoothing (rLOWESS) with a 0.625-second window—half of the MRI sampling rate (TR = 1.25 sec).

To facilitate cross-day and cross-animal comparisons, a z-score transformation was applied. Trial-by-trial baseline correction was performed using a 2-second window preceding the light cue to ensure robust quantification of dopamine excess and account for baseline variability.

### Immunohistochemistry

Brains were harvested immediately following transcardial perfusion and post-fixed in 4% PFA in PBS at 4°C for 24h. Brains were then submerged in 30% sucrose in PBS at 4°C for 18h or until sunk. Brains were frozen in OCT and sectioned with a cryostat into PBS filled wells. 40µm cryosections were mounted onto Superfrost Plus Slides with Fluromount-G with DAPI and imaged.

### Fiber localization and correspondence across mice

To anatomically localize the photometry signal, functional scans were time-averaged for each mouse that underwent dual imaging. In all mice the fiber was visible and the coverage volume was determined by identifying the fiber tip, allowing a tolerance of one voxel in each dimension. This resulted in a coverage volume of approximately 600 µm mediolateral, 400 µm dorsoventral, and 900 µm anteroposterior, enabling estimation of photometry signal spatial correspondence across mice.

---

**Supplementary Figure 1.**
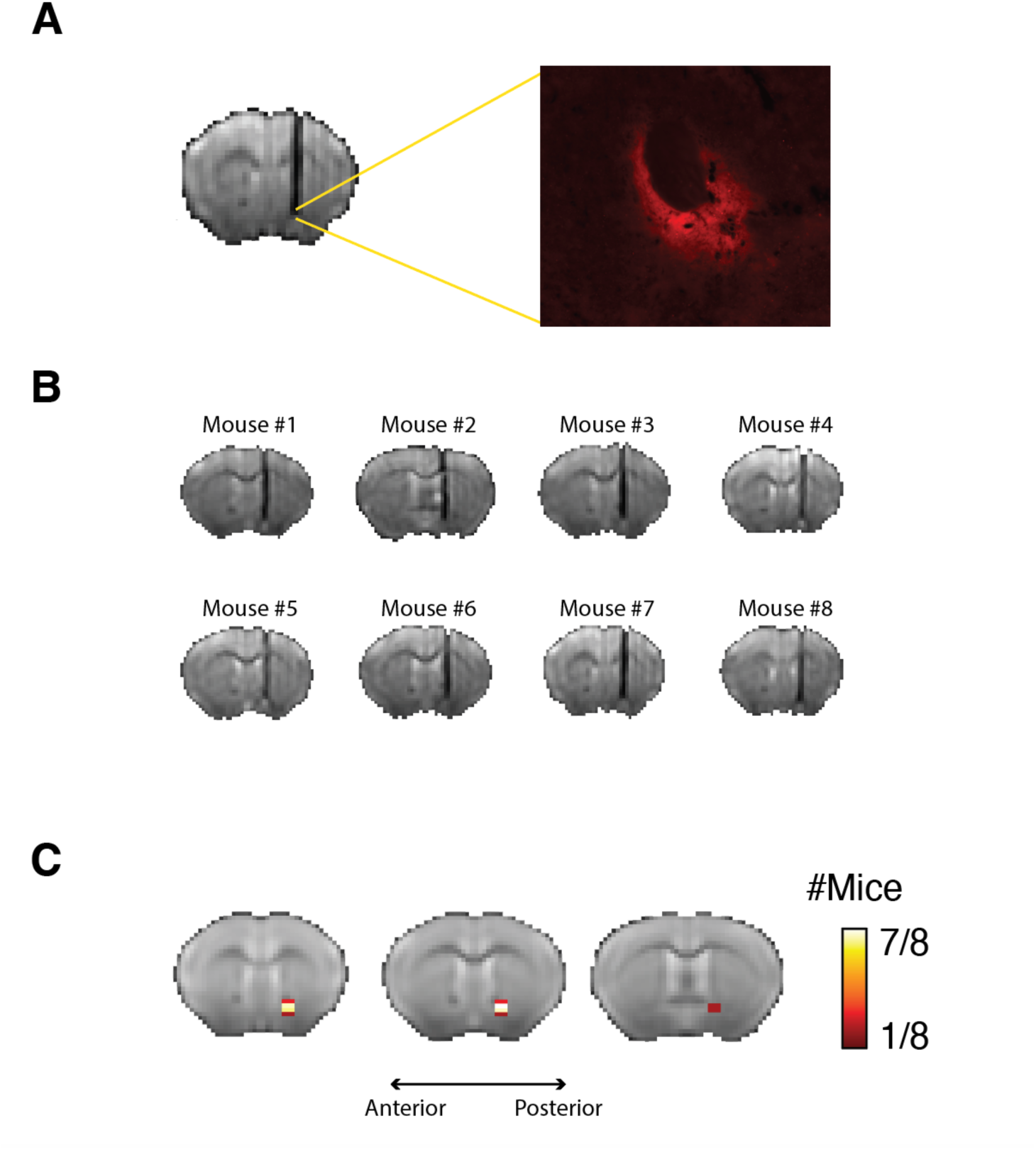
Viral expression and cannula localization. (A) A coronal slice acquired using the functional MRI protocol, showing the placement of the cannula targeting the Nucleus Accumbens (NAc) and the corresponding immunohistochemistry slice from the same region confirming viral expression. All animals included in this experiment showed expression. (B) Cannula placement in each dually imaged mouse in the learning paradigm. (C) Coverage summary of voxels from which GRAB_DA_ signal was acquired across mice, demonstrating consistent targeting of the NAc.

## References

1. Steinberg, E. E., Keiflin, R., Boivin, J. R., Witten, I. B., Deisseroth, K. & Janak, P. H. A causal link between prediction errors, dopamine neurons and learning. Nat Neurosci 16, 966–973 (2013).

2. Day, J. J. & Carelli, R. M. The nucleus accumbens and pavlovian reward learning. Neuroscientist 13 148–159 (2007).

3. Eshel, N., Tian, J., Bukwich, M. & Uchida, N. Dopamine neurons share common response function for reward prediction error. Nat Neurosci 19, 479–486 (2016).

4. Salamone, J. D. & Correa, M. The Mysterious Motivational Functions of Mesolimbic Dopamine. Neuron 76, 470–485 (2012).

5. Wise, R. A. Dopamine, learning and motivation. Nat Rev Neurosci 5, 483–494 (2004).

6. Kutlu, M. G., Zachry, J. E., Melugin, P. R., Tat, J., Cajigas, S., Isiktas, A. U., Patel, D. D., Siciliano, C. A., Schoenbaum, G., Sharpe, M. J. & Calipari, E. S. Dopamine signaling in the nucleus accumbens core mediates latent inhibition. Nat Neurosci 25, 1071–1081 (2022).

7. Waelti, P., Dickinson2, A. & Schultz, W. Dopamine responses comply with basic assumptions of formal learning theory. Nature 412, 43–48 (2001).

8. Berridge, K. C. & Robinson, T. E. Parsing reward. Trends in Neurosciences 26, 507–513 (2003).

9. Roitman, M. F., Wheeler, R. A., Wightman, R. M. & Carelli, R. M. Real-time chemical responses in the nucleus accumbens differentiate rewarding and aversive stimuli. Nat Neurosci 11, 1376– 1377 (2008).

10. M. De Schotten Thiebaut, & Forkel, S.J. The emergent properties of the connected brain. Science 378, 505–510 (2022).

11. Molenberghs, P., Cunnington, R. & Mattingley, J. B. Brain regions with mirror properties: A meta-analysis of 125 human fMRI studies. Neurosci and Biobehav Rev 36, 341–349 (2012).

12. Grill, F., Guitart-Masip, M., Johansson, J., Stiernman, L., Axelsson, J., Nyberg, L. & Rieckmann, A. Dopamine release in human associative striatum during reversal learning. Nat Commun 15(1):59 (2024).

13. Knutson, B., Fong, G. W., Adams, C. M., Varner, J. L. & Hommer, D. Dissociation of reward anticipation and outcome with event-related fMRI. Neuroreport 12, 3683–3687 (2001).

14. Pessiglione, M., Seymour, B., Flandin, G., Dolan, R. J. & Frith, C. D. Dopamine-dependent prediction errors underpin reward-seeking behaviour in humans. Nature 442, 1042–1045 (2006)

15. O’Doherty, J., Dayan, P., Schultz, J., Deichmann, R., Friston, K. & Dolan, R. J. Dissociable roles of ventral and dorsal striatum in instrumental conditioning. Science 304, 452–454 (2004).

16. Haber, S. N. & Knutson, B. The reward circuit: linking primate anatomy and human imaging. Neuropsychopharmacology 35, 4–26 (2010).

17. Daniel, R. & Pollmann, S. A universal role of the ventral striatum in reward-based learning: Evidence from human studies. Neurobiol Learn Mem 114, 90–100 (2014).

18. Shohamy, D. Learning and motivation in the human striatum. Curr Opin Neurobiol 21, 408–414 (2011).

19. Sun, F. et al. A Genetically Encoded Fluorescent Sensor Enables Rapid and Specific Detection of Dopamine in Flies, Fish, and Mice. Cell 174, 481–496.e19 (2018).

20. Amo, R., Matias, S., Yamanaka, A., Tanaka, K. F., Uchida, N. & Watabe-Uchida, M. A gradual temporal shift of dopamine responses mirrors the progression of temporal difference error in machine learning. Nat Neurosci 25, 1082–1092 (2022).

21. Day, J. J., Roitman, M. F., Wightman, R. M. & Carelli, R. M. Associative learning mediates dynamic shifts in dopamine signaling in the nucleus accumbens. Nat Neurosci 10, 1020–1028 (2007).

22. Albin, R. L., Young, A. B. & Penney, J. B. The functional anatomy of basal ganglia disorders. Trends Neurosci 12, 366–375 (1989).

23. Lanciego, J. L., Luquin, N. & Obeso, J. A. Functional neuroanatomy of the basal ganglia. Cold Spring Harb Perspect Med 2, a009621 (2012).

24. Cox, J. & Witten, I. B. Striatal circuits for reward learning and decision-making. Nat Rev Neurosci 20, 482–494 (2019).

25. Sackett, D. A., Saddoris, M. P. & Carelli, R. M. Nucleus accumbens shell dopamine preferentially tracks information related to outcome value of reward. eNeuro 4 (3) ENEURO.0058-17.2017 (2017).

26. Cooper, J. C. & Knutson, B. Valence and salience contribute to nucleus accumbens activation. Neuroimage 39, 538–547 (2008).

27. Ferenczi, E. A. et al. Prefrontal cortical regulation of brainwide circuit dynamics and reward-related behavior. Science 351, aac9698 (2016).

28. Cerri, D. H. et al. Distinct neurochemical influences on fMRI response polarity in the striatum. Nat Commun 15, 1916 (2024).

29. Roitman, M. F., Wheeler, R. A. & Carelli, R. M. Nucleus accumbens neurons are innately tuned for rewarding and aversive taste stimuli, encode their predictors, and are linked to motor output. Neuron 45, 587–597 (2005).

30. Setlow, B., Schoenbaum, G. & Gallagher, M. Encoding predicted outcome and acquired value in orbitofrontal cortex during cue sampling depends upon input from basolateral amygdala. Neuron 39,855–867 (2003).

31. Ray, M. H., Moaddab, M. & McDannald, M. A. Threat and Bidirectional Valence Signaling in the Nucleus Accumbens Core. J Neurosci 42, 817–833 (2022).

32. Otis, J. M., Namboodiri, V. M. K., Matan, A. M., Voets, E. S., Mohorn, E. P., Kosyk, O., McHenry, J. A., Robinson, J. E., Resendez, S. L., Rossi, M. A. & Stuber, G. D. Prefrontal cortex output circuits guide reward seeking through divergent cue encoding. Nature 543, 103–107 (2017).

33. Shen, L., Zhang, G. W., Tao, C., Seo, M. B., Zhang, N. K., Huang, J. J., Zhang, L. I. & Tao, H. W. A bottom-up reward pathway mediated by somatostatin neurons in the medial septum complex underlying appetitive learning. Nat Commun 13, 1194 (2022).

34. Li, N. & Jasanoff, A. Local and global consequences of reward-evoked striatal dopamine release. Nature 580, 239–244 (2020).

35. Lee, H. J., Weitz, A. J., Bernal-Casas, D., Duffy, B. A., Choy, M. K., Kravitz, A. V., Kreitzer, A. C. & Lee, J. H. Activation of Direct and Indirect Pathway Medium Spiny Neurons Drives Distinct Brain-wide Responses. Neuron 91, 412–424 (2016).

36. Long, C., Lee, K., Yang, L., Dafalias, T., Wu, A. K. & Masmanidis, S. C. Constraints on the subsecond modulation of striatal dynamics by physiological dopamine signaling. Nat Neurosci 27, 1977–1986 (2024).

37. Freeze, B. S., Kravitz, A. V., Hammack, N., Berke, J. D. & Kreitzer, A. C. Control of basal ganglia output by direct and indirect pathway projection neurons. J Neurosci 33, 18531–18539 (2013).

38. Kravitz, A. V., Freeze, B. S., Parker, P. R. L., Kay, K., Thwin, M. T., Deisseroth, K. & Kreitzer, A. C. Regulation of parkinsonian motor behaviours by optogenetic control of basal ganglia circuitry. Nature 466, 622–626 (2010).

39. Gallo, E. F., Meszaros, J., Sherman, J. D., Chohan, M. O., Teboul, E., Choi, C. S., Moore, H., Javitch, J. A. & Kellendonk, C. Accumbens dopamine D2 receptors increase motivation by decreasing inhibitory transmission to the ventral pallidum. Nat Commun 9, 1086 (2018).

40. Stauffer, W. R., Lak, A. & Schultz, W. Dopamine reward prediction error responses reflect marginal utility. Curr Biol 24, 2491–2500 (2014).

41. Schultz, W. Dopamine reward prediction-error signalling: a two-component response. Nat Rev Neurosci 17, 183–195 (2016).

42. Fetcho, R. N., Parekh, P. K., Chou, J., Kenwood, M., Chalençon, L., Estrin, D. J., Johnson, M. & Liston, C. A stress-sensitive frontostriatal circuit supporting effortful reward-seeking behavior. Neuron 112, 473–487.e4 (2024).

43. McGuire, J. T. & Botvinick, M. M. Prefrontal cortex, cognitive control, and the registration of decision costs. Proc Natl Acad Sci U S A 107, 7922–7926 (2010).

44. Warden, M. R., Selimbeyoglu, A., Mirzabekov, J. J., Lo, M., Thompson, K. R., Kim, S. Y., Adhikari, A., Tye, K. M., Frank, L. M. & Deisseroth, K. A prefrontal cortex-brainstem neuronal projection that controls response to behavioural challenge. Nature 492, 428–432 (2012).

45. Vertes, R. P. Differential Projections of the Infralimbic and Prelimbic Cortex in the Rat. Synapse 51, 32–58 (2004).

46. Gorgolewski, K. J. et al. BIDS apps: Improving ease of use, accessibility, and reproducibility of neuroimaging data analysis methods. PLoS Comput Biol 13, e1005209 (2017).

47. Chan, R. W., Lee, R. P., Wu, S. Y., Tse, E. L., Xue, Y., Moeller, S. & Chan, K. C. NOise Reduction with DIstribution Corrected (NORDIC) PCA improves signal-to-noise in rodent resting-state and optogenetic functional MRI. in Proc Annu Int Conf IEEE Eng Med Biol Soc 1847–1850 (2022).

48. Desrosiers-Grégoire, G., Devenyi, G. A., Grandjean, J. & Chakravarty, M. M. A standardized image processing and data quality platform for rodent fMRI. Nat Commun 15, 6708 (2024).

49. Pruim, R. H. R., Mennes, M., van Rooij, D., Llera, A., Buitelaar, J. K. & Beckmann, C. F. ICA-AROMA: A robust ICA-based strategy for removing motion artifacts from fMRI data. Neuroimage 112, 267–277 (2015).

